# Experiment-free learning of exoskeleton assistance is not an unsolved problem

**DOI:** 10.64898/2026.06.16.731058

**Authors:** Shuzhen Luo, Menghan Jiang, Sainan Zhang, Junxi Zhu, Shuangyue Yu, Israel Dominguez Silva, Bolei Zhou, Hyunwoo Yuk, Xianlian Zhou, Hao Su

## Abstract

We present three quantitative methods: 1) estimation of exoskeleton mechanical power and energy ratio from published data, 2) a systematic review of the exoskeleton literature on reported energy ratios, and 3) timing correction analysis of the replication experiment, to address concerns raised by Collins et al. (2026) about Luo et al. (2024). Together, these analyses support the reported metabolic reductions and the validity of exoskeleton control via learning in simulation. The critique rests on an unsupported premise: that exoskeleton energy ratios above 4 are physiologically implausible. This premise of Collins et al. (2026) is not supported by the cited evidence, and the error originates in their own cited source. Sawicki and Ferris (2009), the paper they invoke as authority for the limit of 4, state explicitly that "reported values of the muscular efficiency range from 0.10 to 0.34, with many sources assuming an average of ∼0.25." The value of 4 corresponds to this average, it is not a physiological ceiling. Treating an average as a physiological upper limit is a fundamental error. The published exoskeleton literature further contradicts the claim, including work by the authors of the critique themselves (Collins et al., 2015: 4.3; Young et al., 2017: 5.0) and independent work (Malcolm et al., 2013: 4.8; Seo et al., 2017: 6.7). In contrast, our walking energy ratio is 2.4, calculated directly from Fig. 4 of our paper. Our device delivers higher peak torque (14.1 Nm vs. 10.9 Nm, Lim et al., 2019) and achieves a slightly larger metabolic reduction (24.3% vs. 21%). Independent groups have since demonstrated meaningful metabolic reductions using learning-in-simulation frameworks, including Barati et al. (2026, 15.2% mean and 22.5% maximum) and Zhou et al. (2025, ∼20% during running). The claim of Collins et al. (2026) that this problem "remains unsolved" is directly contradicted by these independent results.

The experiment in the critique is not a valid replication of our method. Our controller is a neural network with ∼10,000 parameters learned through deep reinforcement learning in musculoskeletal simulation; the critique instead applies a pre-programmed fixed torque curve with no learnable parameters. Beyond this, the replication contains three methodological errors: 1) a heel-strike timing assumption producing offsets up to 30% of the gait cycle; 2) an averaged torque profile that discards subject-specific control; and 3) a device ∼50% heavier than ours (4.8 kg vs. 3.2 kg) without measuring the metabolic penalty of the added weight. The critique also misreports Samsung data, with reported values approximately double those in the original publication, errors that directly underpin their physiological limit argument.

## Introduction

The core challenges our work addresses are the data scarcity in wearable robotics: training an effective exoskeleton controller requires large amounts of human locomotion data across diverse conditions and individuals, which is expensive and slow to collect experimentally. Most existing learning-based exoskeleton controllers rely on supervised learning from human-collected datasets, which limits scalability and generalization to new users and conditions (Molinaro et al., 2024). The contribution of Luo et al. (2024) is to demonstrate that exoskeleton control policies can be learned in simulation and transferred to hardware, thereby bridging the simulation-to-reality gap. By contrast, deep reinforcement learning in musculoskeletal simulation generates training data through physics-based interaction with a human body model rather than through human experiments, and the resulting policy transfers to real hardware (Park et al., 2026).

Our learned controller is a neural network policy that continuously adapts to each individual’s movement, unlike a hand-designed torque profile with fixed timing and predetermined magnitude. Since publication, multiple groups have developed similar learning-in-simulation approaches for wearable robots. Barati et al. (2026) recently published in IEEE Robotics and Automation Letters, adopt a closely related musculoskeletal simulation and deep reinforcement learning framework for hip exoskeleton control and demonstrate meaningful sim-to-real metabolic reductions on hardware. Related efforts have also appeared from researchers at Carnegie Mellon University (Park et al., 2026; Choi et al., 2026), Tsinghua University (Zuo et al., 2026), Seoul National University (Kim et al., 2025; Leem et al., 2026), and the Chinese Academy of Sciences (Zhou et al., 2025), as summarized in Table 2.

Barati et al. (2026) is a rigorous and well-executed independent study that implemented the same core approach as Luo et al. (2024): musculoskeletal simulation combined with deep reinforcement learning, using a multi-layer perceptron (MLP) policy trained with proximal policy optimization (PPO). Notably, 4 out of 8 subjects in Barati et al. (2026) exhibited energy ratios exceeding 4 (apparent efficiency below 25%), providing direct experimental evidence that an energy ratio of 4 is not a physiological upper limit for exoskeleton-assisted walking.

Collins et al. (2026) assert that exoskeleton energy ratios should not exceed 4 and use this claim as the foundation of their critique. The energy ratio is defined as metabolic savings divided by mechanical work delivered, whether by the exoskeleton or by muscle. Muscular efficiency, also termed apparent efficiency in some studies (e.g., Seo et al., 2016; Barati et al., 2026), measures the same quantity expressed as a fraction rather than a ratio. Energy ratio is the reciprocal of muscular efficiency: an energy ratio of 4 corresponds to a muscular efficiency of 0.25. As Sawicki and Ferris (2009) state directly: “reported values of the muscular efficiency of positive work for mammalian skeletal muscle range from 0.10-0.34, with many sources assuming an average of ∼0.25.” Collins et al. (2026) misstate their own cited source by reporting this average as a strict physiological ceiling while omitting the range the same source provides. On this basis alone, the central premise is unsupported. <u>Treating an average as a physiological upper limit is a fundamental error. Human life expectancy averages approximately 80 years, yet no one would argue that a person who lives to 90 has violated a biological law. Similarly, the average muscular efficiency of ∼0.25 corresponds to an energy ratio of 4, but individuals and conditions that exceed this average are not physiologically impossible, they are simply above average.</u>

The published exoskeleton literature further contradicts the claim of Collins et al. (2026). Although energy ratio is not commonly reported in the exoskeleton literature, when reported it often exceeds 4. Studies by Collins et al. themselves report 4.3 (Collins et al., 2015, Nature), 5.0 (Young et al., 2017), and 5.4 (Ishmael and Lenzi, 2021, Nature Medicine). Additional studies report 4.8 (Malcolm et al., 2013) and up to 6.7 (Seo et al., 2017, cited by Collins et al. themselves). The claimed physiological limit of 4 is contradicted both by the source cited and by the published record, including Collins et al.’s own work.

In contrast, the energy ratios reported in Luo et al. (2024) remain below 4 under both direct calculation from published data (2.4 from Fig. 4 in our paper) and indirect estimation using Collins et al.’s own analytical approach after correcting timing and kinematic errors (3.4, Fig. S3 in this Reply). Under either method, the energy ratios in our paper are well within the range reported in the published exoskeleton literature.

Collins et al. (2026) contain multiple factual errors. For example, they report an assistive torque of 24.1 Nm for the Samsung exoskeleton, whereas the original paper reports 10.9 Nm (Seo et al., 2016) and the device’s hardware torque capability is only 12 Nm. The reported 24.1 Nm therefore exceeds the physical limit of the Samsung robot. Collins et al. (2026) also report apparent-efficiency values that differ materially from the values in the original Samsung papers (see Section 2 for full quantification).

Collins et al. (2026) do not attempt to reproduce the learning-in-simulation methodology proposed in Luo et al. (2024). A meaningful replication would require training an exoskeleton controller in simulation and deploying the learned policy on hardware, as multiple independent groups have since demonstrated (Table 2). Instead, their replication employs a conventional gait-cycle-based controller, a control paradigm widely used in exoskeleton research for more than two decades, applying predefined averaged torque profiles with fixed phase shifts and no controller training in simulation. As a result, they evaluate a fundamentally different control strategy and cannot test the method presented in our paper.

A replication that does not validate its own hardware cannot constitute a meaningful challenge to the method being tested. A valid replication must first demonstrate that the hardware platform can reproduce an established result with a known effective controller, because only then can a failed replication be attributed to the method rather than to the hardware itself. Collins et al. (2026) did not do this. A straightforward validation would have been to implement an established controller, such as the Samsung delayed output feedback controller, on their specific device and confirm a meaningful metabolic reduction. Collins et al. (2026) provide no such validation. Two confounds alone are sufficient to explain their very small metabolic result without implicating our controller: first, their replication device is ∼50% heavier than ours (4.8 kg vs. 3.2 kg), and without a separately measured No Exo baseline, the metabolic penalty of carrying this heavier device cannot be separated from any benefit provided by the controller; second, the incorrect assistance timing, arising from their misinterpretation of our gait phase definition, would substantially reduce delivered mechanical power regardless of which controller was used. Either confound alone could erase an apparent metabolic benefit; together, they make the very small result entirely expected, and entirely uninformative about the validity of our method.

## Methods

### 1) Gait Phase Definition and Timing Analysis

Most existing exoskeleton controllers detect the gait cycle from heel strike, typically measured by sensors on the foot or shank. The controller in Luo et al. (2024) uses a minimal sensor configuration, with one IMU on each thigh and no sensors on the foot or shank, and accordingly does not rely on heel-strike detection. Assistive torque is mapped directly from thigh kinematics to hip joint torque, requiring no gait-phase detection. Fig. 3b in in Luo et al. (2024) is a post-processing visualization used for analysis, not the timing reference used by the real-time controller.

**Fig. 1.**
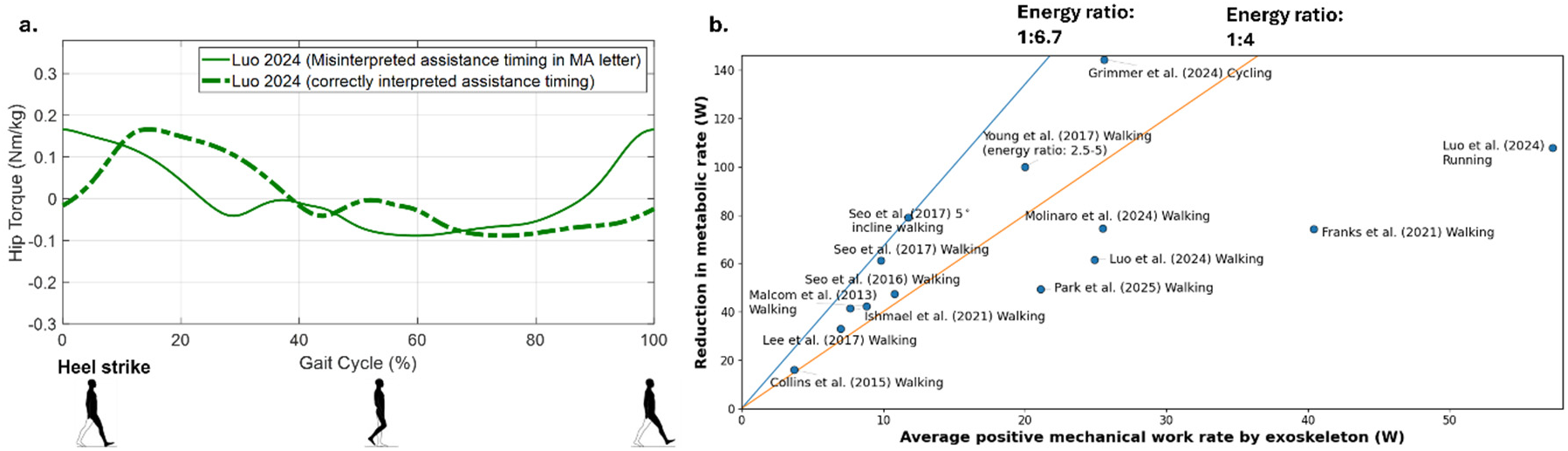
Misinterpreted assistance torque timing in Collins et al. (2026) vs. correctly interpreted timing in Luo et al. (2024). (a) Collins et al. (2026) assume that 0% gait phase in Fig. 3b of Luo et al. (2024) corresponds to heel strike. In contrast, in our paper 0% gait phase is defined as the peak of assistive torque, which can occur up to 30% of the gait cycle after heel strike. The dashed line illustrates a representative 18% timing alignment used to demonstrate the effect of correcting this misinterpretation for power estimation; it does not represent a fixed correction that can replace participant-specific assistance timing. (b) Prior studies report energy ratios exceeding 4 (orange line), with values reaching 6.7 (summarized in Fig. 2 of this Reply). Many studies do not report energy ratios, including Slade and Collins (2022), Sartori (Durandau et al., 2022), Gregg (Divekar et al., 2024), and Huang and Si (Tu et al., 2021).

In Luo et al. (2024), 0% gait phase is defined by the learned assistance profile itself, aligned to the peak of assistive torque rather than to heel strike. Collins et al. (2026) misread this convention, assuming that 0% gait phase in Fig. 3b corresponds to heel strike. Collins et al. (2026) is incorrect: thigh IMUs cannot detect heel strike, so heel strike is neither measured nor used in control. The peak of assistive torque can occur substantially after heel strike, with prior hip exoskeleton studies reporting timing differences of up to approximately 30% of the gait cycle (Young et al., 2017). The heel-strike assumption therefore introduces a systematic timing offset that distorts downstream estimates of exoskeleton power, energy ratio, and the phase offsets used in the replication experiment. Applying fixed 5%, 10%, or 15% phase shifts to an averaged torque profile does not correct this error; it only tests several open-loop profiles around the wrong reference frame. This is a key reason why the replication by Collins et al. (2026) fails to reproduce our result.

### 2) Our Reported Metabolic Reductions Are Physiologically Plausible and Consistent with Prior Work

Luo et al. (2024) paper already provides enough torque and power information to check the approximate walking power scale. From the walking segment around 1.25 m/s in Fig. 4 of Luo et al. (2024), the estimated exoskeleton positive mechanical power is approximately 24.9 W, giving an energy ratio of ∼2.4 using the reported walking metabolic reduction. This estimate is not presented as a new full-cohort power measurement; rather, it shows that the appropriate walking power scale is around 24 W, not the much lower value of 11.2 ± 2.0 W estimated by Collins et al. (2026). Their lower estimate arises from incorrect timing assumptions in the power calculation. Even when using Collins et al. (2026)’s own estimation framework, correcting the timing and gait-kinematics assumptions yields an estimated exoskeleton positive power of 17.9 W, which still gives an energy ratio below 4. Thus, their conclusion depends on substantially underestimating the exoskeleton positive mechanical power.

Barati et al. (2026) reports a mean reduction of 15.2% and a maximum individual reduction of 22.5% with a mean peak torque of 9.8 Nm (up to 10.5 Nm) and 17.3 W mean positive power (primarily flexion power assistance), demonstrating that the learning-in-simulation paradigm generalizes to an independently developed system at a different institution. The larger metabolic reduction in Luo et al. (2024) is mechanistically expected: our controller delivers higher total mechanical power, on the order of ∼24 W, by assisting both hip flexion and extension throughout the gait cycle, whereas Barati et al.’s device delivers power primarily during hip flexion. Combined with a larger peak torque (14.1 Nm vs. 9.8 Nm mean), the broader gait-cycle coverage and higher mechanical output directly account for the larger metabolic benefit. Notably, Barati et al.’s maximum individual reduction of 22.5% with smaller peak torque and smaller exoskeleton power already approaches our mean of 24.3%, further confirming that reductions in this range are physiologically achievable. If our metabolic result were anomalous or our power estimate inflated, it would break this cross-device consistency, yet it does not.

**Fig. 2.**
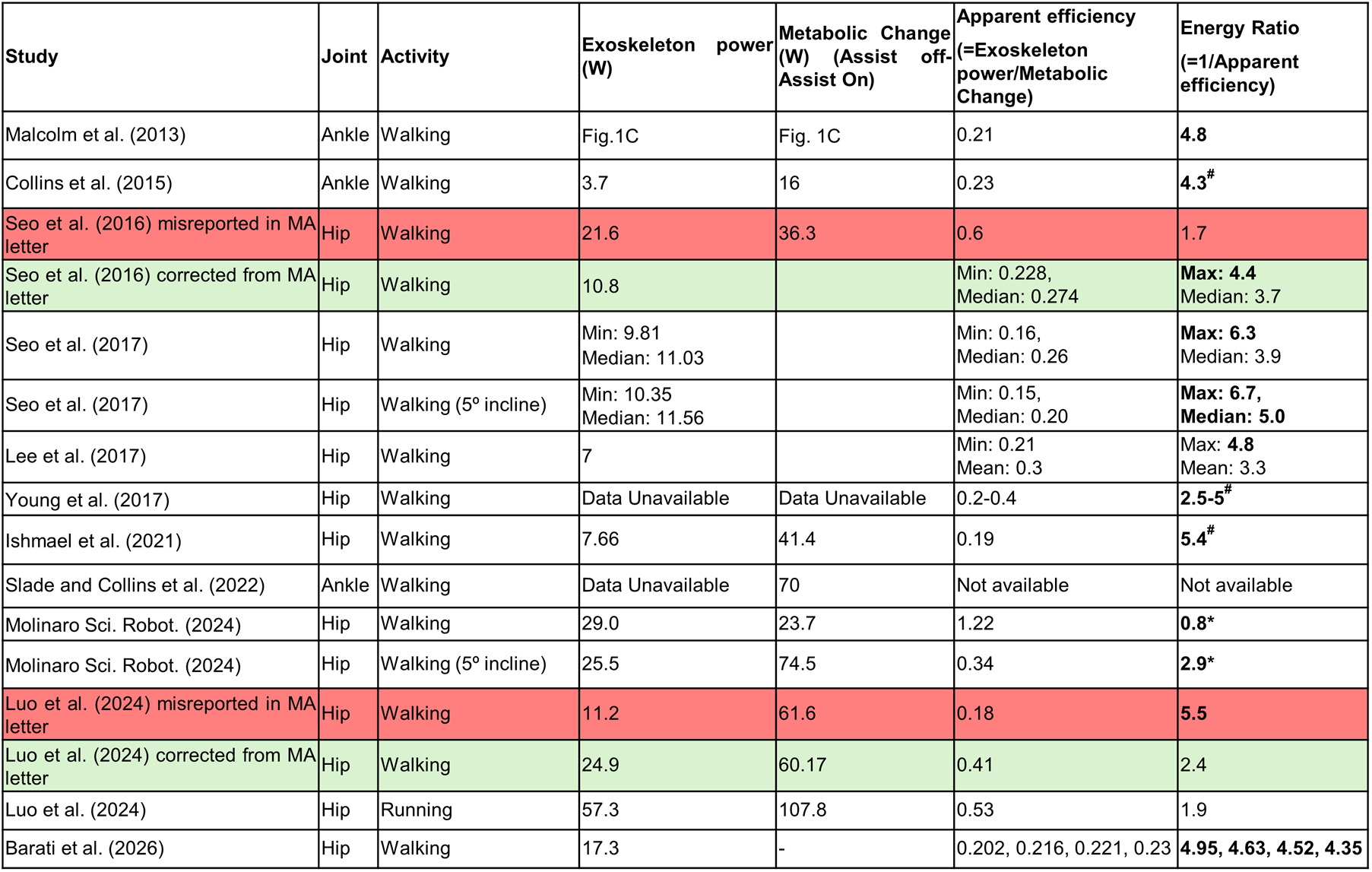
Although energy ratio is not typically reported in the exoskeleton literature, multiple studies that do report it show values exceeding 4, including work by the authors of Collins et al. (2026) themselves (Collins et al., 2015: 4.3; Young et al., 2017: 5.0; Ishmael and Lenzi, 2021: 5.4) and other studies (Malcolm et al., 2013: 4.8; Seo et al., 2017: up to 6.7). Energy ratios exceeding 4 are common in prior exoskeleton studies including papers by multiple authors of Collins et al. (2026) (marked as #) and correctly interpreted values place Luo et al. (2024) well below this range. When corrected values are used, the energy ratios for Luo et al. (2024) remain well below 4 for walking and running and do not approach any physiological limit. Energy ratio is highly sensitive, e.g., for two closely related activities using the same controller, Molinaro et al. reports an energy ratio of 0.8 during level walking (presented in Collins et al. (2026)) and 2.9 during 5° incline walking (based on published data), over a threefold difference. Furthermore, <u>Barati et al. (2026) reported that 4 out of 8 subjects exhibited energy ratios exceeding 4 (apparent efficiency below 25%), providing direct experimental evidence that an energy ratio of 4 is not a physiological limit for exoskeleton-assisted walking.</u>

**Table 1.** Four independently developed hip exoskeletons show a consistent torque-to-power-to-metabolic relationship. The 24% reduction in our study is squarely within this published trend, directly refuting Collins et al. (2026)’s "physiologically implausible" claim.

| Study | Device mass (kg) | Applied peak torque (Nm) | Bilateral mean positive power (W) | Mean metabolic reduction (%) |
| --- | --- | --- | --- | --- |
| Lim et al., 2019 (Samsung) | 2.1 | 10.9 | ~20 | 19.8 (mean), max 24.9 (max) |
| Lee et al., 2017 (Samsung) | 2.4 | 10.9 | ~20 | 21 |
| Barati et al., RAL 2026 | 2.9 | 9.8 | 16.9-17.3 | 15.2 (mean), 22.5 (max) |
| Luo et al. (2024) | 3.2 | 14 | ~24.9 | ~24 |

The mechanical basis for our results is also consistent with prior soft exosuit studies. Single direction hip assistance alone can produce meaningful metabolic reductions: Tricomi et al. (2024) reported a 17.79% reduction in young adults during uphill walking using flexion-only assistance and a 10.48% reduction in older adults during level ground walking. Kim et al. (2022) reported a 14.8% reduction using a hip flexion-only exosuit; and Kim et al. (2019) reported a 9.3% reduction using hip-extension assistance during treadmill walking at 1.5 m/s. Our device assisted both hip flexion and hip extension, with timing aligned to phases of positive hip motion. Because mechanical power is torque multiplied by angular velocity, assistive torque delivered during both major hip work phases can produce substantial positive mechanical power. Therefore, a metabolic reduction in the 20% range is mechanically and physiologically reasonable when sufficient positive power is delivered with appropriate timing.

Collins et al. (2026) contains multiple factual errors, including two factual errors regarding Samsung exoskeleton data. Both errors incorrectly favor Collins et al. (2026)’s "physiological limit" claim.

i. Peak torque. Collins et al. (2026) states the Samsung peak torque as 24.1 Nm. The original Samsung publication reports a maximum of 10.9 Nm (Seo et al., 2016). The 24.1 Nm in Collins et al. (2026) does not appear in the Samsung literature, and exceeds the 12 Nm hardware capability of the Samsung device as shown in Supplementary Fig. S1. It cannot have come from the published record.
ii. Energy ratio. Collins et al. (2026)’s Table S1 reports an apparent efficiency of 0.6 for Seo et al., implying an energy ratio of only 1.7 (= 1/0.6). The actual published energy ratio reaches 6.7 (= 1/0.15) for individual subjects in Seo et al. (2017, ICORR). Samsung’s own published data therefore exceeds Collins et al. (2026)’s claimed "ratio ≤ 4" limit.

These errors fail on two levels. First, even with the correct Samsung values, Samsung’s own data already exceed Collins et al. (2026)’s "ratio ≤ 4" limit (ratio up to 6.7); the limit does not hold in the very literature Collins et al. (2026) cites as authority. Second, our walking energy ratio is 2.4, far below 4 regardless of whether such a limit exists. Neither Collins et al. (2026)’s argument nor its errors bear on the validity of Luo et al. (2024).

Collins et al. (2026) Fig. S2 also misreports assistive torque, depicting 24.1 Nm peak torque, more than double 10.9 Nm reported in the original study. It then incorrectly claims that our assistive torque is substantially smaller. In fact, our peak torque (14.1 Nm) exceeds that (10.9 Nm) reported by Seo et al. (2016) while achieving slightly larger metabolic reductions (24.3%) than those (21%) reported in a follow-up study from the same group (Lee et al., 2017). These errors in Collins et al. (2026) are not minor as they materially affect Collins et al. (2026)’s analysis and the conclusions drawn from it.

Collins et al. (2026)’s analysis of the stair-climbing energy ratio is invalid. It combines metabolic data from stairmill walking with exoskeleton power estimates from outdoor stair climbing, which involve different stair heights and cadences, rendering Collins et al. (2026)’s reported energy ratio value meaningless. The stair-climbing metabolic cost reduction reported in Luo et al. (2024) is an experimentally measured outcome; the absence of concurrent exoskeleton power measurement means an energy ratio cannot be derived, not that the metabolic result is unverifiable.

### 3) Methodological Differences Between the Replication Study and the Original Method

The experiment by Collins et al. (2026) differs from Luo et al. (2024) in several fundamental ways. The original method uses a neural network controller trained via deep reinforcement learning in simulation; the alternative experiment applies a pre-programmed fixed torque curve with no learnable parameters. These are fundamentally different control architectures, and timing adjustment or averaging correction cannot bridge this gap.

The so-called "replication" combines several confounds that are each sufficient to erase an apparent metabolic benefit: incorrect timing, an averaged rather than participant-specific torque profile, and a substantially heavier device. Metabolic rate measurements are inherently noisy, e.g., in Collins et al. (2015), the signal-to-noise ratio was only 2.8, meaning noise was approximately 36% of the measured effect, compared to ∼1000 for wearable IMU joint-angle signals. A very small metabolic result obtained under wrong timing, averaged control, and added device mass is therefore not strong evidence against Luo et al. (2024); it is the expected outcome of a noisy and confounded experiment.

First, the replication relies on a heel-strike-based timing assumption that contradicts the phase definition in Luo et al. (2024) and misaligns assistive torque with hip angular velocity. Collins et al. (2026)’s fixed 5%, 10%, and 15% shifts do not solve this problem because they still shift an averaged open-loop profile around the wrong heel-strike reference frame.

Second, Collins et al. (2026) applies a single averaged torque profile across all participants, whereas our controller delivers participant-specific assistance. Averaging removes the individualized timing and magnitude learned by the controller and reduces a state-feedback policy to a generic open-loop profile. A fixed average profile is not expected to reproduce the benefit of a controller fitted to each participant.

Third, the replication exoskeleton in Collins et al. (2026) was 50% heavier than ours (4.8 kg vs. 3.2 kg). An extra 1.6 kg worn on the body is large enough to change the metabolic outcome of a walking experiment, particularly when the mass is carried near the hip.

Unlike a point mass carried at the waist, the exoskeleton mass is distributed along the thighs and moves with each stride, increasing rotational inertia of the legs and amplifying the metabolic cost of swinging the limbs. Prior load-carriage data suggest this additional mass imposes a substantial metabolic penalty, with larger effects for lighter participants (Browning et al., 2007). Critically, Collins et al. (2026) did not separately measure the metabolic cost of wearing the heavier device without assistance, an Assist Off versus No Exo comparison. Without that measurement, the controller benefit cannot be separated from the weight penalty. The numbers make this concrete: if assistance reduces metabolic rate by 16% relative to Assist Off, but wearing the heavier device adds a 15% penalty relative to No Exo, the net result appears as only a 1% reduction, essentially what Collins et al. (2026) reports. This scenario is plausible given the device mass, and Collins et al. (2026) cannot rule it out because it was never measured.

Taken together, the so-called replication experiments by Collins et al. (2026) tests a fundamentally different control strategy and device configuration from those in Luo et al. (2024), and therefore does not constitute a valid assessment of our findings.

### 4) Musculoskeletal Model and Learning-in-Simulation Framework Are Reproducible Within Their Stated Scope

Learning in simulation is widely adopted in robotics, and Luo et al. (2024) demonstrate its applicability to exoskeleton control. In robotics, reproducibility refers to independent implementation of a methodology and its successful application to related systems, rather than exact numerical replication across hardware platforms. The evidence above satisfies this standard.

Collins et al. (2026) characterizes the learning-in-simulation framework in Luo et al. (2024) as insufficiently described and not reproducible. This is directly contradicted by the broad independent uptake of the methodology since publication. Barati et al. (2026), published in IEEE Robotics and Automation Letters, trained a deep-RL hip exoskeleton controller in musculoskeletal simulation and reported metabolic reductions up to 22.5% individual (15.2% mean). Park et al. (2026, CMU) deployed a predictive musculoskeletal simulation pipeline on a hip exoskeleton and stated that "learning entirely in simulation is feasible." Zhou et al. (2025, Chinese Academy of Sciences) reported ∼20% metabolic reduction during running in a follow-up paper. Kim et al. (2025, Seoul National University) deployed an end-to-end deep-RL controller on a lower-limb exoskeleton. The same framework has been extended to full lower-limb rehabilitation exoskeletons for individuals with neuromuscular impairments using a teacher-student policy distillation paradigm in simulation (Luo et al., 2026). Additional independent implementations of the learning-in-simulation approach appear across Tsinghua University, NJIT, Rutgers, NYU, and the open-source MyoAssist platform (Table 2).

**Table 2.** Publications on learning in simulation for versatile control of exoskeletons.

| Venue | Paper | Institution | Method and Results |
| --- | --- | --- | --- |
| 2026<br>RAL | End-to-end Policy Learning for Hip exoskeleton Control via Deep RL and Reflex-based Musculoskeletal Simulation | KIST, Gyeongsang National University | sim-to-real RL on a hip exoskeleton; 15.2% mean metabolic reduction, up to 22.5% individual, at 9.8 Nm peak torque |
| 2025<br>ROBIO | A Simulation-based Lower-limb Exoskeleton Control Method | Chinese Academy of Sciences | Simulation-based hip exoskeleton control; follow-up reports ~20% metabolic reduction during running |
| 2025<br>IROS | A Deep Reinforcement Learning based End-to-End Control Framework for Lower Limb Exoskeletons with Smooth Movement Transitions | Seoul National University | End-to-end DRL for smooth locomotion transitions across multiple activities |
| 2026<br>ICLR | Exo-plore: Exploring Exoskeleton Control Space through Human-Aligned Simulation | Seoul National U; Northeastern U | Published in top machine learning venue; validates simulation-based exoskeleton assistance optimization |
| 2026 | SMAT: Staged Multi-Agent Training for Co-Adaptive Exoskeleton Control | NJIT and collaborators | Extends learning-in-simulation to multi-agent human-exoskeleton co-adaptation |
| 2026 | Musculoskeletal Motion Imitation for Learning Personalized Exoskeleton Control Policy in Impaired Gait | CMU | Extends musculoskeletal RL toward personalized exoskeleton control for impaired-gait populations |
| 2026 | Learning Hip Exoskeleton Control Policy via Predictive Neuromusculoskeletal Simulation | CMU | Musculoskeletal simulation, RL, and policy distillation; deployed on a hip exoskeleton |
| 2026<br>arXiv | Embodied Human Simulation for Quantitative Design and Analysis of Interactive Robotics | Tsinghua University | Reports 8 hours GPU training for comparable musculoskeletal control, refuting Collins et al. (2026)'s 240-hour comparison |
| 2026<br>JNER | Robust neural network controller for a lower limb exoskeleton with minimal sensor configuration: a deep reinforcement learning approach with policy distillation | ERAU, NJIT, NYU | Extends sim2real to a full lower-limb exoskeleton for individuals with neuromuscular impairments via simulation validation |
| 2026<br>Biorob | Gait asymmetry from unilateral weakness and improvement with ankle assistance: a reinforcement learning based simulation study | NJIT, Rutgers | RL-based simulation study demonstrating ankle exoskeleton assistance reduces gait asymmetry caused by unilateral weakness |
| 2026<br>Biorob | Reinforcement-Learning-Based Assistance Reduces Squat Effort with a Modular Hip-Knee Exoskeleton | NJIT | Applies RL in simulation to optimize hip-knee exoskeleton assistance for squatting |
| Open-source platform | MyoAssist: open-source Python toolkit for musculoskeletal control via deep RL | Facebook AI; Northeastern University | Open-source musculoskeletal RL toolkit for wearable robots, used in NeurIPS MyoChallenge |

Collins et al. (2026) claims that the musculoskeletal model in Luo et al. (2024) is unspecified, that the network architecture and training procedures lack key details, and that convergence criteria are absent. Luo et al. (2024) explicitly states that the musculoskeletal model is derived from the established MASS (Muscle-Actuated Skeletal System) framework (Lee et al., 2019), with minor modifications. The learning framework, including network architectures, reward functions, and training procedures, is fully specified; Collins et al. (2026)’s claim that these elements are insufficiently described is therefore unsupported. Convergence was assessed following standard practice in deep reinforcement learning.

### Pre-submission correspondence

Collins et al. (2026) discuss pre-submission correspondence in their public preprint. For completeness, we clarify the record. In 2024, Dr. Collins requested metabolic-rate data, which we shared. Collins et al. subsequently submitted their Matters Arising to Nature in December 2025 and posted it publicly on bioRxiv in April 2026 while the Nature review process was ongoing. Before submitting and publicly posting the critique, Collins et al. did not raise with us the specific technical issues that now form the central basis of their critique, including the energy ratio calculation, gait phase definition, Samsung comparison, or replication protocol, for clarification, correction, or discussion. Thus, these technical claims were not previously presented to us prior to publication of the critique. The characterization of our position as a refusal to share is inconsistent with the documented correspondence and release timeline. We provide this context only to clarify the sequence of events; the scientific issues are addressed in the technical sections above.

Collins et al. (2026) state that pre-submission outreach was led by Dr. Robert Gregg. For the record, neither the first author nor the corresponding author received any communication from Dr. Robert Gregg.

## Supplementary Materials

Extensive supplementary materials supporting the findings and reproducibility of Luo et al. (2024) have been released for independent verification, including executable code, raw experimental data, and simulation files. These Supplementary Materials provide detailed clarification and supporting analyses corresponding to the sections in Collins et al. (2026) Supplementary Materials, documenting relevant definitions, calculations, and methodological distinctions using data reported in the original paper and prior literature.

**Supplementary figure S1.**
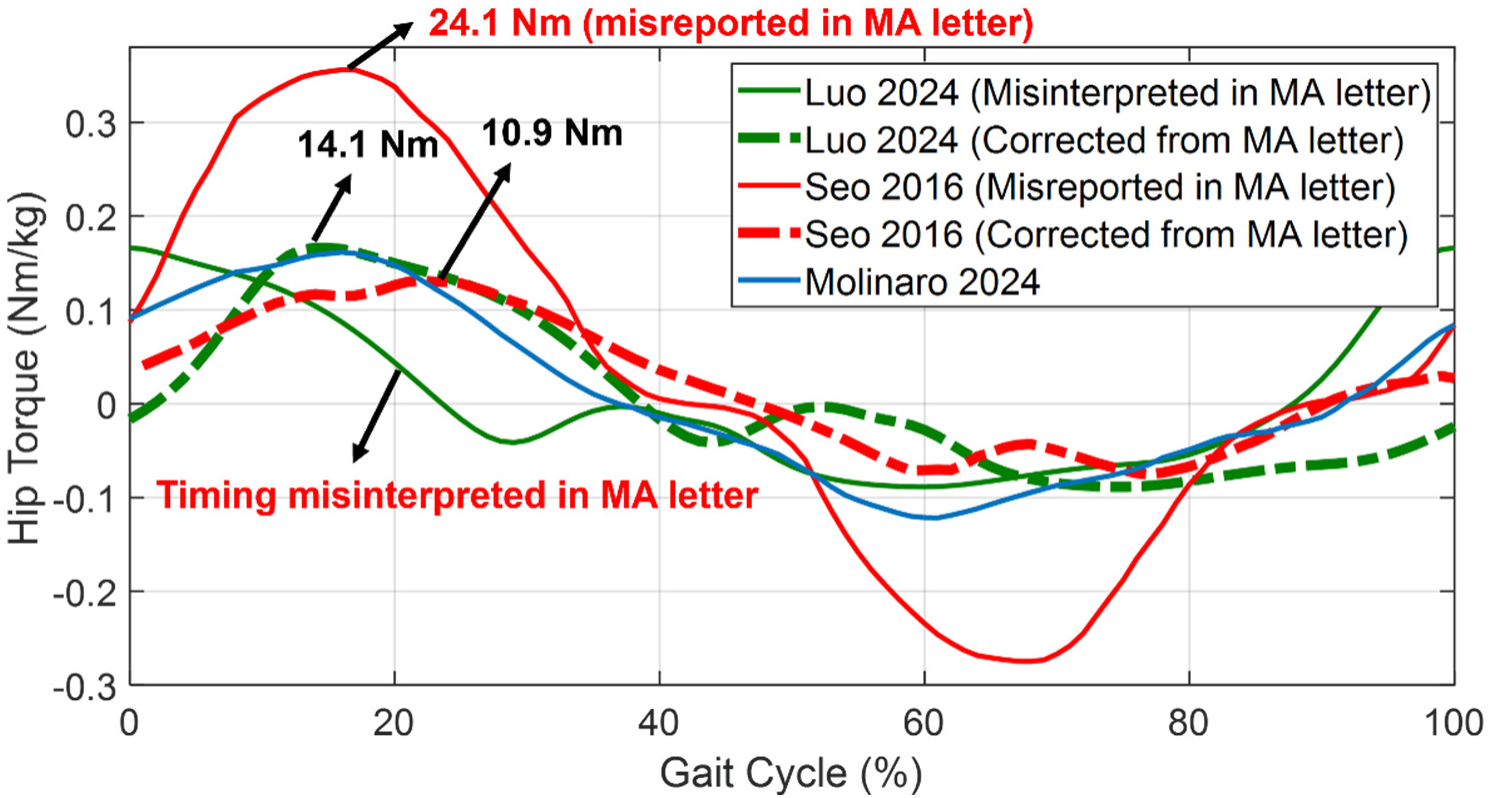
Collins et al. (2026) contains two errors that materially undermine its critique. First, the peak assistive torque reported for the Samsung exoskeleton (Seo et al., 2016) is 10.9 Nm, whereas Collins et al. (2026) incorrectly reports 24.1 Nm. Second, Collins et al. (2026) assumes that the peak assistive torque in Luo et al. (2024) occurs at heel strike (0% gait cycle), whereas in our study the peak torque occurs typically after heel strike, with 0% gait phase defined at the torque peak. Luo et al. (2024) report a peak torque of 14.1 Nm with a ∼24.3% metabolic reduction, which is physiologically meaningful and consistent with follow-up studies from the same group, including Lim et al. (2019) (10.9 Nm, ∼19.8% reduction) and Lee et al. (2017) (10.9 Nm, ∼21% reduction). These errors in Collins et al. (2026) distort cross-study comparisons and directly underpin its proposed “physiological limit” argument.

### S1) Assistance Timing is Misinterpreted in Collins et al. (2026) Resulting Incorrect Exoskeleton Power and Metabolic Energy Cost

As detailed in Main Text Section 1 and Fig. 1, Collins et al. (2026) treats 0% assistance phase as heel strike, although Luo et al. (2024) defined 0% by the peak of assistive torque. This timing error is consequential: Lee et al. (2017) showed that even a 4% gait-cycle timing offset reduced metabolic benefit from 21.1% to 14.7%, and an 8% offset reduced it further to 13.6-14.7%. Thus, Collins et al. (2026)’s heel-strike-referenced timing and fixed phase shifts cannot validate or refute a participant-specific learned controller.

Under both direct calculation from Fig. 4 of Luo et al. (2024) (2.4) and Collins et al. (2026)’s own analytical method with corrected timing and kinematics (3.4, Fig. S3), our walking energy ratio remains below 4.

**Supplementary figure S2.**
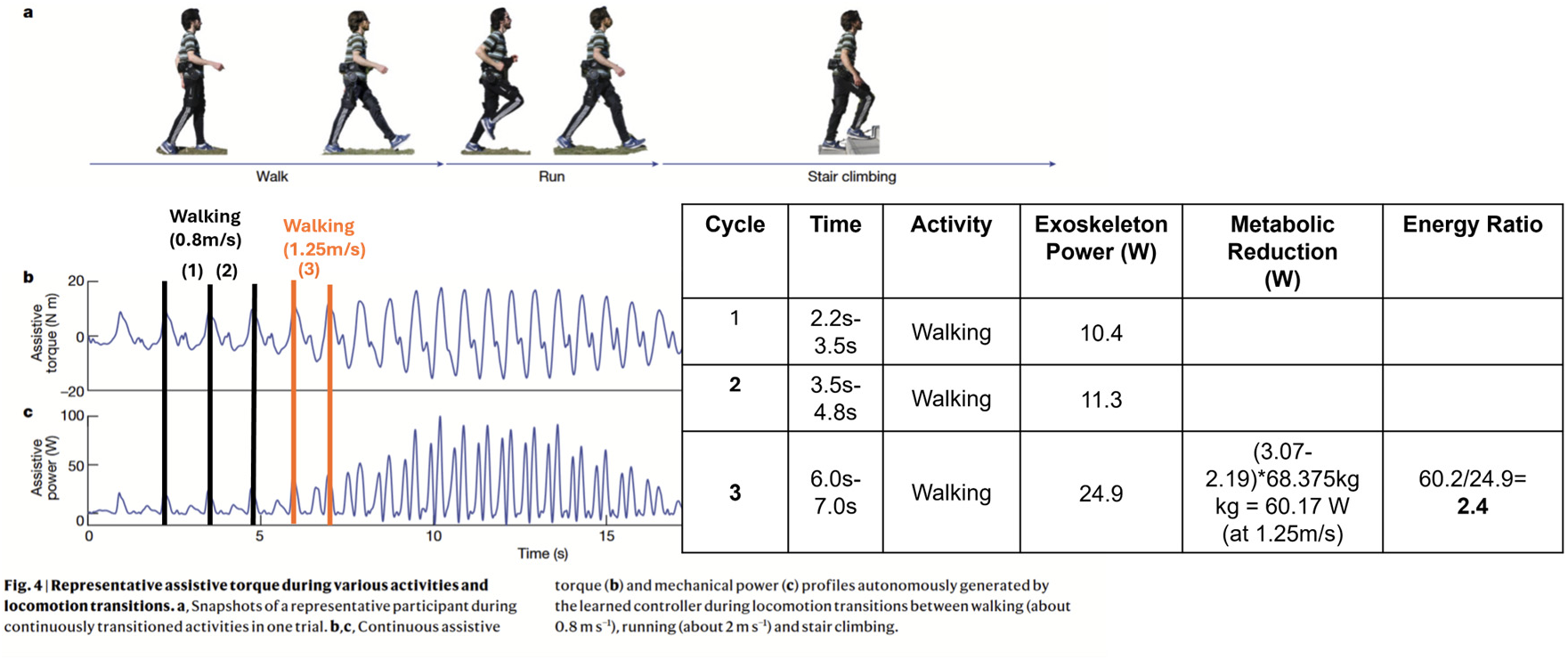
Energy ratio calculation derived directly from Fig. 4 of Luo et al. (2024) yields an energy ratio below 4. Exoskeleton torque and power during walking and the transition to running show that the energy ratios remain below 4. Luo et al. (2024) reported metabolic cost measurements at 1.25 m/s walking and 2.0 m/s running. At 1.25 m/s walking, the exoskeleton positive power derived from Fig. 4 of Luo et al. (2024) is 24.9 W, whereas Collins et al. (2026) estimates only 11.2 W. Collins et al. (2026) estimate is comparable to the power at 0.8 m/s walking (11.3 W) and is therefore not representative of the 1.25 m/s walking condition used for the metabolic analysis. Using the published power values, the walking energy ratio is 2.4.

Collins et al. (2026) estimate also assumes negligible kinematic effects of assistance, which is contradicted by published measurements showing assistance modifies joint angle and velocity profiles. Under either calculation, our walking energy ratio is below 4.

Collins et al. (2026) underestimates exoskeleton power during 1.25 m/s walking, reporting 11.2 W, essentially the same as the 11.3 W reported at 0.8 m/s (Fig. S2). This indicates Collins et al. (2026) estimation does not scale with walking speed. Three factors contribute: heel-strike-referenced timing causing power-velocity misalignment, neglect of joint kinematic changes under assistance, and use of an averaged rather than participant-specific torque profile.

Published exoskeleton studies, including by the authors of Collins et al. (2026), exceed Collins et al. (2026)’s claimed limit of 4 (full list in Main Text Section 1 and Fig. 2). Collins et al. (2026)’s premise is contradicted by its own cited authority.

**Supplementary figure S3.**
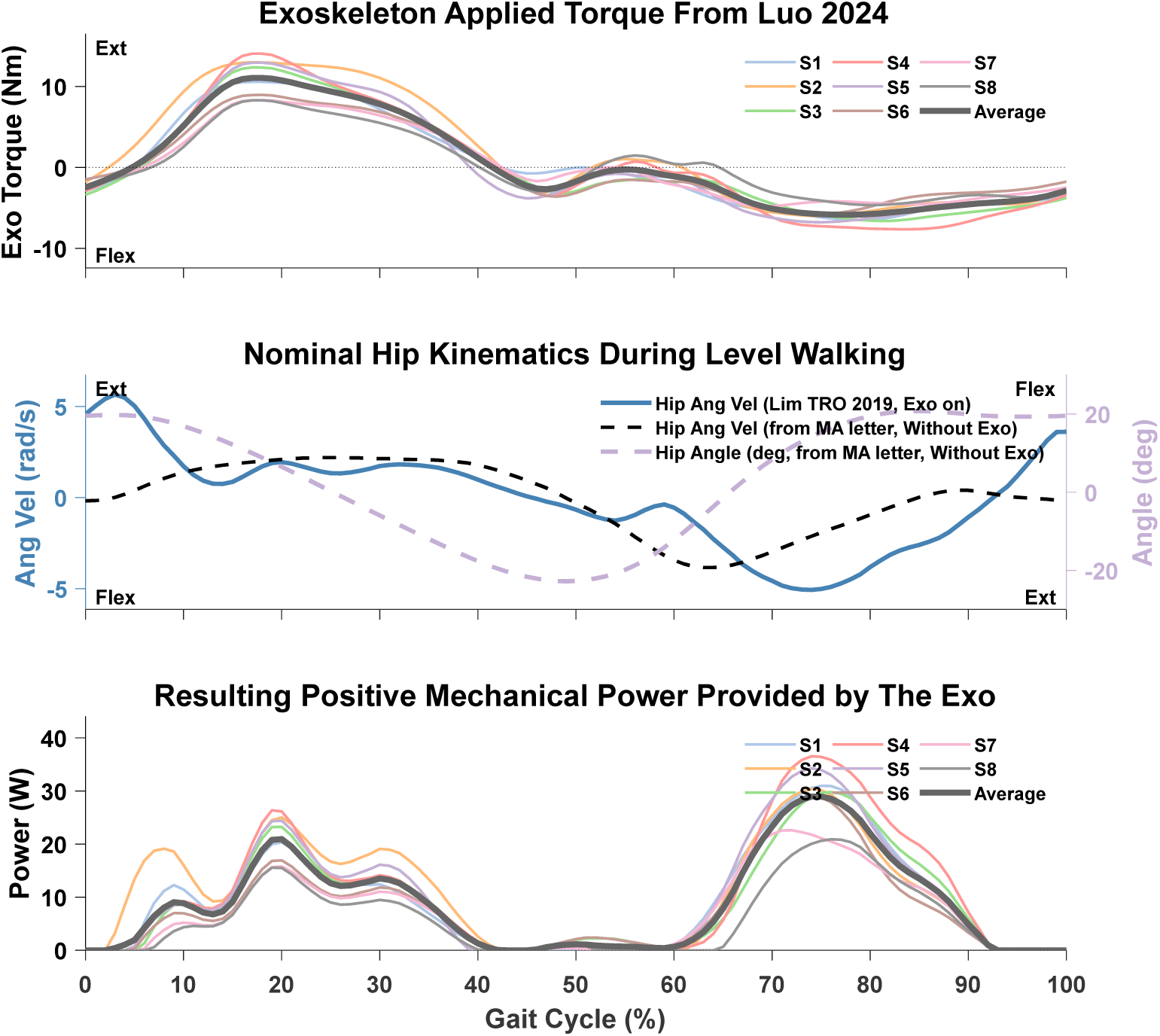
Using Collins et al. (2026)’s replication code and procedure, applying an 18% timing correction as a representative mid-range value within the up-to-30% offset reported by Young et al. (2017), and scaling hip angular velocity to account for walking speed differences (4 km/h = 1.11 m/s in Lim et al., 2019 vs. 1.25 m/s in Luo et al., 2024), yields an estimated exoskeleton power of 17.9 W and an energy ratio of ∼3.4 based on hip kinematics from Lim et al. (2019). Using the data directly reported in Luo et al. (2024), the energy ratio during walking at 1.25 m/s is ∼2.4 (Supplementary Fig. S2). Thus, whether using Collins et al. (2026)’s estimation approach or our directly reported measurements, the energy ratio remains well below the claimed physiological limit of 4.

### S2) Collins et al. (2026) Experiment Fails to Test the Reported Method Due to Timing, Averaging, And Device Mass Differences

The energy ratio estimated using Collins et al. (2026)’s own method is approximately 3.4 (Fig. S3), while direct calculation from Fig. 4 of Luo et al. (2024) yields 2.4, both below the claimed limit of 4. The replication fails on three grounds detailed in Main Text Section 3. First, it assumes a heel-strike-referenced gait cycle, whereas 0% gait phase in Luo et al. (2024) corresponds to the peak of assistive torque, which can occur up to ∼30% of the gait cycle after heel strike (Young et al., 2017). This timing mismatch substantially reduces delivered mechanical power. Second, it applies a single averaged torque profile across all participants, discarding the participant-specific timing and magnitude learned by our controller; simple phase shifts of an averaged profile cannot recover individualized assistance (Lee et al., 2017). Third, the device was ∼50% heavier than ours (4.8 kg vs. 3.2 kg) without a No Exo baseline, as detailed above.

### S3) Learning-In-Simulation Framework Is Reproducible Within Stated Scope and Standard Practice of Robotics Research

In robotics, reproducibility is established when independent groups implement the same methodology and achieve consistent outcomes on related systems, rather than through exact numerical replication across hardware platforms. Table 2 and Main Text Section 4 document broad independent uptake of the learning-in-simulation framework across research groups worldwide. In particular, Barati et al. (2026) applied the same musculoskeletal simulation method using approximately 20 muscles (compared to 208 in our work) and reported meaningful metabolic reductions, demonstrating that the core framework generalizes across different levels of model complexity.

Collins et al. (2026)’s concern about absence of a reported random seed reflects a low-level implementation detail rarely specified in journal publications. Our training uses standard PPO initialization and the outcome is consistent across multiple seeds.

Convergence in Luo et al. (2024) was assessed following standard practice in reinforcement learning: stabilization of the training reward curve, with convergence reached when the reward changed by less than 5% across 10 consecutive validation evaluations.

To clarify the policy evaluation procedures questioned in Collins et al. (2026), trained control policies were evaluated using multiple performance metrics, including gait stability, assistive torque smoothness, assistance timing, symmetry, and assistance power. For the GetExoReward and GetHumanReward functions referenced by the authors of Collins et al. (2026), the corresponding formulations are provided in Eqs. (7-9) and Eqs. (7-9, 13, 14) of the Methods section in Luo et al. (2024).

For completeness regarding the simulation implementation, we clarify how the relative motion between the human body segments and the exoskeleton was computed. The relative translation was first computed in the global frame and then transformed into the corresponding local body frame before evaluating the bushing force. This is a standard coordinate-transformation procedure for computing attachment forces between coupled rigid bodies. The translational stiffness coefficients kx, ky, and kz and damping coefficients cx, cy, and cz have units of N/m and N·s/m, respectively. The rotational stiffness coefficients αx, αy, and αz and damping coefficients βx, βy, and βz have units of N·m/rad and N·m·s/rad, respectively. These implementation details are now explicitly clarified here and are also provided in the released extended codebase for verification.

**Supplementary figure S4.**
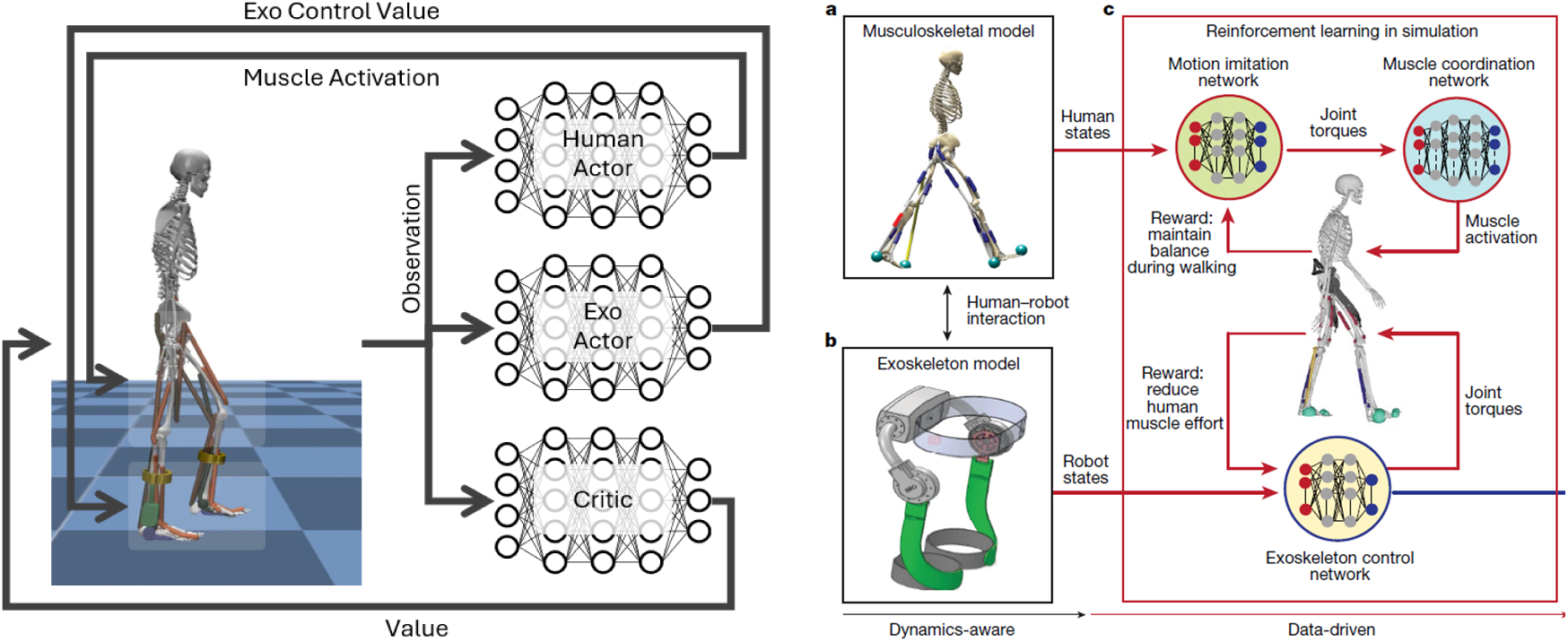
(Left) MyoAssist platform based on MyoSuite have very similar learning in simulation framework as our paper (right) for wearable robot control and this approach is becoming readily available in the machine learning community as annual MyoChallenge in the Conference on Neural Information Processing Systems (NeurIPS).

The publicly available MyoSuite and MyoAssist platforms, based on Facebook AI Research, adopt a learning-in-simulation approach closely aligned with our paper for both exoskeleton and prosthesis control. MyoAssist is featured in the annual MyoChallenge at the Conference on Neural Information Processing Systems, demonstrating that learning-in-simulation for musculoskeletal control has become standard practice in the machine-learning community. The SMAT (Staged Multi-Agent Training for Co-Adaptive Exoskeleton Control) paper by Yuan et al. (2026), currently under review, is based on the MyoAssist platform and states the intent to open source the implementation once published. In addition, Exo-plore by Leem et al. (2026) from Seoul National University further illustrates this trend: this work presents an open-source neuromechanical simulation framework that combines musculoskeletal simulation with deep reinforcement learning to optimize hip exoskeleton control parameters without requiring real human experiments.

Collins et al. (2026)’s critique that our method lacks a metabolic model or nervous system is not relevant to the validity of our approach. Our Hill-type musculoskeletal model with force-length and force-velocity dynamics provides the physiological substrate, and the neural-network actor functions as motor command generation.

Musculoskeletal human models are comparable to humanoid robots, and learning-based control methods developed for one extend naturally to the other. Our 50-DOF biomechanical human model coupled to a 2-DOF hip exoskeleton is within the standard complexity of contemporary humanoid-robot control.

### S4) Musculoskeletal Model Is Specified and Adopted from An Established Framework

Collins et al. (2026)’s claim that our musculoskeletal model is unspecified is incorrect. As stated in Luo et al. (2024), the model is adopted from the established MASS framework (Lee et al., 2019), which provides full descriptions of muscle parameters, body segments, joint definitions, contact dynamics, and integration with deep RL.

### S5) Modeling and Computational Efficiency Comparison Support Our Methods

Multiple critiques in Collins et al. (2026) appear to arise from misunderstandings of the deep reinforcement learning rather than from replication attempts. For instance, Collins et al. (2026) claims that our neural network lacks activation functions and output distributions (p. 19), despite these being explicitly defined in the published pseudocode. As a result, multiple comments focus on peripheral implementation details rather than the core deep learning framework.

Collins et al. (2026) raises the ∼8 hour training time reported in Luo et al. (2024) as suspicious, comparing it to the ∼240 hour training time in Simos et al. (2025) This is directly contradicted by the recent Tsinghua study on embodied human simulation for exoskeleton co-design, which independently reports ∼8 hours of GPU training for a comparable musculoskeletal control task (Zuo et al., 2026). Collins et al. (2026)’s own letter further acknowledges that the difference from Simos et al. (2025) "likely relates to the much larger neural networks required" in that study, meaning the two studies operate at fundamentally different model scales and are not comparable. Collins et al. (2026)’s comparison is therefore not a valid critique.

More broadly, Collins et al. (2026)’s reliance on the “240 hours of training” reported in Simos et al. (2025) reflects a misunderstanding of how training time scales in deep learning and how different learning paradigms should be compared. Luo et al. (2024) adopt the same human walking control network and muscle coordination network as the established MASS model (Lee et al., 2019), augmented only with an additional exoskeleton control network. This exoskeleton module is substantially simpler than the human controller, with much smaller input and output dimensions, and introduces only a few additional degrees of freedom corresponding to bilateral hip-assistance torques. As a result, the added computational burden per simulation step is limited, and the overall training time remains comparable to that of human-only walking control, approximately 8 hours. <u>This estimate is further supported by recent work from Tsinghua University, where Zuo et al. (2026) reported that training a robust musculoskeletal human control policy within an embodied human simulation framework required approximately 8 hours, followed by a substantially shorter optimization stage for interactive robot design and control</u>.

### S6) Simulation Artifacts Reflect Contact Dynamics and Do Not Indicate Physical Implausibility

Supplementary Movie 1 in Luo et al. (2024) is a post-edited compilation of several separately rendered simulation clips, created only to illustrate the learning-in-simulation concept. <u>The video is a conceptual demonstration video for visualization purposes, not a real-time simulation output, and is not the same as the quantitative simulation data and results reported in our paper. The video shows outdoor activities for illustration purposes, which do not correspond to our training scenarios. The actual musculoskeletal simulation used in the paper contains 208 muscle-tendon units based on MASS framework (Lee et al., 2019, ACM SIGGRAPH); rendering all 208 muscles simultaneously in a video would visually overcrowd the figure and make the body itself hard to see, so for visual clarity the movie shows only a small subset of muscles, and their color does not change as they activate. The video therefore cannot be used to read off muscle activity.</u>

Collins et al. (2026) identified a very brief foot-contact force in one frame of the supplementary simulation videos that appears to violate the friction constraint and interpreted this as evidence of physical implausibility. However, such transient irregularities are well-known numerical optimization artifacts (Stewart and Trinkle, 1996, Werling et al. 2021) in rigid-body physics simulation with intermittent contact, particularly during rapid contact transitions such as heel strike and toe-off.

Our simulations use MASS platform (Lee et al., 2019), which is built on the DART (Dynamic Animation and Robotics Toolkit) simulation engine (Lee et al., 2018). DART handles contacts and collisions using an implicit time stepping, velocity-based linear complementarity problem (LCP) formulation with an approximated Coulomb friction cone represented using a polyhedral approximation (Lee et al., 2018; Stewart and Trinkle, 1996). As acknowledged in the DART documentation itself, the framework uses approximate Coulomb friction cone conditions rather than an exact nonlinear friction formulation. These formulations are widely used because of their computational efficiency and robustness, but they are also known to produce occasional non-smooth or transient contact-force artifacts, especially during discontinuous contact transitions and under finite solver iterations.

The interpretation that the brief contact-force irregularities are numerical simulation artifacts is also supported by subsequent work built directly on DART. Werling et al. (2021), in the Nimble/DART differentiable physics work, describe DART’s contact handling as a boxed LCP formulation using the Dantzig solver inherited from ODE. The authors explicitly note that boxed LCPs are not theoretically guaranteed to be solvable in all cases, but are common in practice because of their computational speed and high-quality empirical results. Their discussion further highlights the practical trade-offs inherent in rigid-body contact simulation with friction. These contact-modeling limitations are not unique to DART but are common across major physics engines. Comparative simulator studies have shown that contact and friction modeling remain open challenges across major physics engines. Erez et al. (2015), in a comparison of Bullet, Havok, MuJoCo, ODE, and PhysX, emphasized that contact dynamics with friction are computationally difficult and that practical simulators necessarily rely on tractable approximations and numerical regularization strategies. Their benchmark results showed that each engine tends to perform best on the type of system it was designed for, with differing trade-offs in contact handling across simulators, particularly in contact-rich tasks and under larger integration time steps. Similarly, Le Lidec et al. (2024) reviewed modern contact models in robotics and noted that simulation discrepancies can arise from contact-model assumptions, solver accuracy, friction-cone linearization, and time discretization effects. They specifically discuss that polyhedral approximations of the friction cone can bias tangential friction forces and alter simulated trajectories relative to exact Coulomb friction formulations. The review also notes that underdetermined contact configurations are common in robotics applications, including legged locomotion, and can generate spurious internal tangential forces or contact residuals depending on the numerical solver and convergence conditions. Their analysis further demonstrates that different contact formulations can produce similar system-level motions despite differing levels of complementarity or friction residuals.

The difficulty of obtaining reliable contact and friction behavior across simulators is further illustrated by Taylor, Drumwright, and Hsu (2016), who compared four open-source physics engines on a robotic grasping task and identified multiple simulator-specific failure modes, including constraint drift, truncation error from early termination of iterative solvers, and jitter from correction of interpenetration, that can produce unexpected contact force behavior even in well-established simulation frameworks. Crucially, their analysis highlights that non-convergent iterative solves and numerical artifacts are recognized failure modes of LCP-based rigid-body solvers, not indicators of invalid underlying physics.

Therefore, the relevant question is not whether every instantaneous contact-force sample perfectly satisfies an ideal continuous-time Coulomb friction cone, but whether any deviations are rare, bounded, localized to contact transitions, and non-influential to the reported system-level behavior. In our simulations, the observed friction-cone residuals were isolated, short-duration events occurring during contact transition frames rather than sustained stance phases. They did not produce persistent slipping, nonphysical propulsion, unstable gait behavior, or systematic alteration of the reported experimental conclusions. Consequently, these transient residuals are more appropriately interpreted as known numerical artifacts of rigid-body contact simulation rather than evidence of invalid overall dynamics.

We provide a new supplementary video of walking simulation showing both muscle activations and ground reaction forces, including curve plots of joint angles, ground reaction forces, and exoskeleton assistance torque (https://www.youtube.com/watch?v=0CHMQ6bModE). The video shows an unedited, continuous simulation output during walking. Unlike the original conceptual movie, this video corresponds to the quantitative simulation workflow and confirms that the actual simulations run smoothly and continuously across multiple consecutive gait cycles.

## Conclusion

The central premise of Collins et al. (2026) is not supported by the cited evidence: they treat an average value as a physiological upper limit. Sawicki and Ferris (2009), the very source Collins et al. (2026) cite as authority, state explicitly that "reported values of the muscular efficiency (η_muscle) of positive work for mammalian skeletal muscle range from 0.10 to 0.34, with many sources assuming an average of ∼0.25." Because energy ratio is the reciprocal of muscular efficiency, the average efficiency of ∼0.25 corresponds to an energy ratio of 4, the mean of the reported range, not a physiological ceiling. From the exoskeleton literature, the claim is similarly unsupported. The authors of Collins et al. (2026) themselves have published energy ratios exceeding 4 in their own prior work: Collins et al. (2015) report 4.3, Young et al. (2017) report 5.0, and Ishmael and Lenzi (2021) report 5.4. Additional independent studies report 4.8 (Malcolm et al., 2013) and up to 6.7 (Seo et al., 2017). Most directly, Barati et al. (2026) report that 4 out of 8 subjects exceeded an energy ratio of 4 under identical conditions, bilateral hip exoskeleton, level-ground walking, with individual values of 4.95, 4.63, 4.52, and 4.35. These empirical observations directly falsify the claim that 4 is a physiological upper limit: if it were, these values could not have been measured. In contrast, our own energy ratio, calculated directly from Fig. 4 of Luo et al. (2024), is 2.4, based on ∼24.9 W exoskeleton positive mechanical power during walking at 1.25 m/s, well below 4 under direct calculation, and still below 4 (at ∼3.4, corresponding to 17.9 W) even when using Collins et al.’s own analytical framework after correcting their timing error.

The replication by Collins et al. (2026) is invalid on four independent grounds: the controller type is fundamentally different, a pre-programmed torque curve with no learnable parameters rather than a neural network trained in simulation; the timing reference was incorrect; the replication applied an averaged open-loop torque profile rather than reproducing the learning-in-simulation pipeline central to Luo et al. (2024); and the replication device was approximately 50% heavier than ours (4.8 kg vs. 3.2 kg) without separately measuring the metabolic penalty of carrying the heavier device. When consistent definitions and appropriate experimental distinctions are applied, the results reported in Luo et al. (2024) remain internally consistent, physiologically plausible, and aligned with prior state-of-the-art findings. The conclusions of Collins et al. (2026) are not supported by the analyses presented. Their multiple factual errors, including misreported prior study values and the misrepresentation of an average as a physiological limit, are not minor oversights; publishing such errors carries a meaningful risk of misleading the field and setting back progress in wearable robotics.

## Notes

https://www.biorxiv.org/content/10.64898/2026.04.01.715109v1

### Competing Interest Statement

The authors have declared no competing interest.

## References

Luo, S., Jiang, M., Zhang, S., Zhu, J., Yu, S., Dominguez Silva, I., Wang, T., Rouse, E., Zhou, B., Yuk, H., Zhou, X. and Su, H. Experiment-free exoskeleton assistance via learning in simulation. Nature 630, 353–359 (2024).

Seo, K., Hyung, S., Kim, J. and Shim, Y. Fully autonomous hip exoskeleton saves metabolic cost of walking. In Proc. IEEE International Conference on Robotics and Automation 4628–4635 (2016).

Barati H, Kim S, Nguyen TX, Lee J, Park YJ., End-to-end Policy Learning for Hip exoskeleton Control via Deep Reinforcement Learning and Reflex-based Musculoskeletal Simulation. IEEE Robotics and Automation Letters, 2026. https://ieeexplore-custom.ieee.org/document/11543257, DOI: 10.1109/LRA.2026.3699246.

Barati H, Kim S, Nguyen TX, Lee J, Park YJ. Deep reinforcement learning for hip exoskeleton control via predictive simulation of reflex-based human gait. IEEE International Conference on Robotics and Automation (ICRA), 2026.

Seo, K., Lee, J. and Park, Y. J. Autonomous hip exoskeleton saves metabolic cost of walking uphill. In Proc. IEEE International Conference on Rehabilitation Robotics 246–251 (2017).

Collins, S. H., De Groote, F., Gregg, R. D., Huang, H., Lenzi, T., Sartori, M., Sawicki, G. S., Si, J., Slade, P. and Young, A. J. Experiment-free learning of exoskeleton assistance remains an unsolved problem. bioRxiv (2026). 10.64898/2026.04.01.715109

Collins, S. H., Wiggin, M. B. and Sawicki, G. S. Reducing the energy cost of human walking using an unpowered exoskeleton. Nature 522, 212–215 (2015).

Slade, P., Kochenderfer, M. J., Delp, S. L. and Collins, S. H. Personalizing exoskeleton assistance while walking in the real world. Nature 610, 277–282 (2022).

Lim, B., Kim, Y., Jung, J., Seo, K., Ryu, J. and Shim, Y. Delayed output feedback control for gait assistance with a robotic hip exoskeleton. IEEE Transactions on Robotics 35, 1055–1062 (2019).

Zhang, J., Fiers, P., Witte, K. A., Jackson, R. W., Poggensee, K. L., Atkeson, C. G. and Collins, S. H. Human-in-the-loop optimization of exoskeleton assistance during walking. Science 356, 1280–1284 (2017).

Lee, S., Lee, K., Park, M. and Lee, J. Scalable muscle-actuated human simulation and control. ACM Transactions on Graphics 38, 1–13 (2019).

Lee, J., Hwangbo, J., Wellhausen, L., Koltun, V. and Hutter, M. Learning quadrupedal locomotion over challenging terrain. Science Robotics 5, eabc5986 (2020).

Lee, J., Seo, K., Lim, B., Jang, J., Kim, K. and Choi, H. Effects of assistance timing on metabolic cost, assistance power, and gait parameters for a hip-type exoskeleton. In Proc. IEEE International Conference on Rehabilitation Robotics 498–504 (2017).

Peng, X. B., Andrychowicz, M., Zaremba, W. and Abbeel, P. Sim-to-real transfer of robotic control with dynamics randomization. In Proc. IEEE International Conference on Robotics and Automation 3803–3810 (2018).

Sawicki, G. S. and Ferris, D. P. Powered ankle exoskeletons reveal the metabolic cost of plantar flexor mechanical work during walking with longer steps at constant step frequency. Journal of Experimental Biology 212, 21–31 (2009).

Li, M., Wen, Y., Gao, X., Si, J. and Huang, H. Toward expedited impedance tuning of a robotic prosthesis by reinforcement learning. IEEE Transactions on Robotics 38, 407–420 (2022).

Stewart, D. E. and Trinkle, J. C. An implicit time-stepping scheme for rigid body dynamics with inelastic collisions and Coulomb friction. International Journal for Numerical Methods in Engineering 39, 2673–2691 (1996).

Erez, T., Tassa, Y., and Todorov, E. (2015). "Simulation Tools for Model-Based Robotics: Comparison of Bullet, Havok, MuJoCo, ODE, and PhysX." Proceedings of the IEEE International Conference on Robotics and Automation (ICRA), Seattle, WA, USA, pp. 4397–4404. DOI: 10.1109/ICRA.2015.7139807.

Le Lidec, Q., Jallet, W., Montaut, L., Laptev, I., Schmid, C. and Carpentier, J. Contact models in robotics: a comparative analysis. IEEE Transactions on Robotics (2024).

Lee, J., Grey, M. X., Ha, S., Kunz, T., Jain, S., Ye, Y., Srinivasa, S. S., Stilman, M., and Liu, C. K. (2018). "DART: Dynamic Animation and Robotics Toolkit." Journal of Open Source Software, 3(22):500.

Taylor, J. R., Drumwright, E. M., and Hsu, J. (2016). "Analysis of Grasping Failures in Multi-Rigid Body Simulations." Proceedings of the IEEE International Conference on Simulation, Modeling, and Programming for Autonomous Robots (SIMPAR), San Francisco, CA, USA, pp. 295–301.

Werling, K., Omens, D., Lee, J., Exarchos, I., and Liu, C. K. (2021). "Fast and Feature-Complete Differentiable Physics Engine for Articulated Rigid Bodies with Contact Constraints." Proceedings of Robotics: Science and Systems (RSS)

Ma, Y., Wei, C., Zuo, C., Zhang, C. and Sui, Y. Bipedal balance control with whole-body musculoskeletal standing and falling simulations. In Proc. Conference on Robot Learning 305, 4641–4656 (2025).

Simos, M., Chiappa, A. S. and Mathis, A. Reinforcement learning-based motion imitation for physiologically plausible musculoskeletal motor control. arXiv arXiv:2503.14637 (2025).

Kim, J., Heimgartner, R., Lee, G., Karavas, N., Nathanson, D., Galiana, I. and Menard, N. Reducing the metabolic rate of walking and running with a versatile, portable exosuit. Science 365, 668–672 (2019).

Young, A., Foss, J., Gannon, H. and Ferris, D. P. Influence of power delivery timing on the energetics and biomechanics of humans wearing a hip exoskeleton. Frontiers in Bioengineering and Biotechnology 5, 4 (2017).

Durandau, G., Rampeltshammer, W. F., van der Kooij, H. and Sartori, M. Neuromechanical model-based adaptive control of bilateral ankle exoskeletons: biological joint torque and electromyogram reduction across walking conditions. IEEE Transactions on Robotics 38, 1380–1394 (2022).

Tu, X., Li, M., Liu, M., Si, J. and Huang, H. H. A data-driven reinforcement learning solution framework for optimal and adaptive personalization of a hip exoskeleton. In Proc. IEEE International Conference on Robotics and Automation (2021).

Ishmael, M. K., Archangeli, D. and Lenzi, T. Powered hip exoskeleton improves walking economy in individuals with above-knee amputation. Nature Medicine 27, 1783–1788 (2021).

Malcolm, P., Derave, W., Galle, S. and De Clercq, D. A simple exoskeleton that assists plantarflexion can reduce the metabolic cost of human walking. PLoS ONE 8, e56137 (2013).

Divekar, N. V., Thomas, G. C., Yerva, A. R., Frame, H. B. and Gregg, R. D. A versatile knee exoskeleton mitigates quadriceps fatigue in lifting, lowering, and carrying tasks. Science Robotics 9, eadr8282 (2024).

Ferris, D. P., Czerniecki, J. M. and Hannaford, B. An ankle-foot orthosis powered by artificial pneumatic muscles. Journal of Applied Biomechanics 21, 189–197 (2005).

Molinaro, D. D., Kang, I. and Young, A. J. Estimating human joint moments unifies exoskeleton control, reducing user effort. Science Robotics 9, eadi8852 (2024).

Park, J., Min, S., Chang, P. S., Lee, J., Park, M. S. and Lee, J. Generative GaitNet. In ACM SIGGRAPH 2022 Conference Proceedings 1–9 (2022).

Kim, M., Baek, W. J. and Park, J. A deep reinforcement learning based end-to-end control framework for lower limb exoskeletons with smooth movement transitions. In Proc. IEEE/RSJ International Conference on Intelligent Robots and Systems 12005–12012 (2025).

Zhou, Y., Chen, C., Zhang, J., Liu, Y., Zhang, R., Zhang, Z. and Wu, X. A simulation-based lower-limb exoskeleton control method. In Proc. IEEE International Conference on Robotics and Biomimetics 1453–1458 (2025).

Leem, G., Lee, J., Lee, J., Song, S. and Won, J. Exo-plore: exploring exoskeleton control space through human-aligned simulation. In Proc. International Conference on Learning Representations (2026).

Yuan, Y., Androwis, G. and Zhou, X. SMAT: staged multi-agent training for co-adaptive exoskeleton control. arXiv arXiv:2603.07618 (2026).

Choi, I., Park, I., Halilaj, E. and Kang, I. Musculoskeletal motion imitation for learning personalized exoskeleton control policy in impaired gait. arXiv arXiv:2604.09431 (2026).

Park, I., Song, C. and Kang, I. Learning hip exoskeleton control policy via predictive neuromusculoskeletal simulation. arXiv arXiv:2603.04166 (2026).

Zuo, C., Xu, J., Vergnolle, M. Q. and Sui, Y. Embodied human simulation for quantitative design and analysis of interactive robotics. arXiv arXiv:2603.09218 (2026).

NeuMove Lab. MyoAssist: an open-source Python toolkit for simulating and optimizing assistive devices in neuromechanical simulations. Documentation and GitHub repository (2025).

MyoHub. MyoSuite: a collection of musculoskeletal environments and tasks simulated with the MuJoCo physics engine for machine-learning-based biomechanical control. Documentation and GitHub repository (2022–2026).

Browning, R. C., Modica, J. R., Kram, R. and Goswami, A. The effects of adding mass to the legs on the energetics and biomechanics of walking. Medicine & Science in Sports & Exercise 39, 515–525 (2007).

MyoChallenge. Annual MyoChallenge at the Conference on Neural Information Processing Systems. Challenge documentation and website.

Tricomi E, Missiroli F, Xiloyannis M, Lotti N, Zhang X, Stefanakis M, Theisen M, Bauer J, Becker C, Masia L. Soft robotic shorts improve outdoor walking efficiency in older adults. Nature Machine Intelligence. 2024 Oct;6(10):1145–55.

Kim J, Quinlivan BT, Deprey LA, Arumukhom Revi D, Eckert-Erdheim A, Murphy P, Orzel D, Walsh CJ. Reducing the energy cost of walking with low assistance levels through optimized hip flexion assistance from a soft exosuit. Scientific reports. 2022 Jun 29;12(1):11004.

Yuan, Y., Androwis, G. and Zhou, X. Gait asymmetry from unilateral weakness and improvement with ankle assistance: a reinforcement learning based simulation study. 2026 IEEE International Conference on Biomedical Robotics and Biomechatronics (BioRob) (2026). arXiv:2602.18862.

Ratnakumar, N., Tohfafarosh, M.H., Jauhri, S. and Zhou, X. Reinforcement-learning-based assistance reduces squat effort with a modular hip-knee exoskeleton. 2026 IEEE International Conference on Biomedical Robotics and Biomechatronics (BioRob) (2026). arXiv:2602.17794.

